# Longitudinal analysis of humoral immunity against SARS-CoV-2 Spike in convalescent individuals up to 8 months post-symptom onset

**DOI:** 10.1101/2021.01.25.428097

**Authors:** Sai Priya Anand, Jérémie Prévost, Manon Nayrac, Guillaume Beaudoin-Bussières, Mehdi Benlarbi, Romain Gasser, Nathalie Brassard, Annemarie Laumaea, Shang Yu Gong, Catherine Bourassa, Elsa Brunet-Ratnasingham, Halima Medjahed, Gabrielle Gendron-Lepage, Guillaume Goyette, Laurie Gokool, Chantal Morrisseau, Philippe Bégin, Valérie Martel-Laferrière, Cécile Tremblay, Jonathan Richard, Renée Bazin, Ralf Duerr, Daniel E. Kaufmann, Andrés Finzi

## Abstract

Functional and lasting immune responses to the novel coronavirus (SARS-CoV-2) are currently under intense investigation as antibody titers in plasma have been shown to decline during convalescence. Since the absence of antibodies does not equate to absence of immune memory, we sought to determine the presence of SARS-CoV-2-specific memory B cells in COVID-19 convalescent patients. In this study, we report on the evolution of the overall humoral immune responses on 101 blood samples obtained from 32 COVID-19 convalescent patients between 16 and 233 days post-symptom onset. Our observations indicate that anti-Spike and anti-RBD IgM in plasma decay rapidly, whereas the reduction of IgG is less prominent. Neutralizing activity in convalescent plasma declines rapidly compared to Fc-effector functions. Concomitantly, the frequencies of RBD-specific IgM+ B cells wane significantly when compared to RBD-specific IgG+ B cells, which increase over time, and the number of IgG+ memory B cells which remain stable thereafter for up to 8 months after symptoms onset. With the recent approval of highly effective vaccines for COVID-19, data on the persistence of immune responses are of central importance. Even though overall circulating SARS-CoV-2 Spike-specific antibodies contract over time during convalescence, we demonstrate that RBD-specific B cells increase and persist up to 8 months post symptom onset. We also observe modest increases in RBD-specific IgG+ memory B cells and importantly, detectable IgG and sustained Fc-effector activity in plasma over the 8-month period. Our results add to the current understanding of immune memory following SARS-CoV-2 infection, which is critical for the prevention of secondary infections, vaccine efficacy and herd immunity against COVID-19.

## Introduction

Severe acute respiratory syndrome coronavirus-2 (SARS-CoV-2), the causative agent of the ongoing Coronavirus disease 2019 (COVID-19) pandemic, is highly contagious and has infected close to a 100 million people worldwide and caused over 2 million deaths since its discovery. The dynamics and persistence of immune responses in individuals infected with SARS-CoV-2 is currently under needful investigation. Several studies with acute and convalescent COVID-19 patients have showed prompt induction of B and T cell responses upon infection, along with the detection of antigen-specific memory B and T cell responses several weeks into convalescence (1-6). Additionally, antibodies induced upon infection have been shown to protect from SARS-CoV-2 reinfection in animal models (7-9). Passive immunization using neutralizing monoclonal antibody treatments decreased viral loads in animal studies and in patients with COVID-19 (10, 11). The viral target of neutralizing antibodies is the highly immunogenic trimeric Spike (S) glycoprotein, which facilitates SARS-CoV-2 entry into host cells via its receptor-binding domain (RBD) that interacts with angiotensin-converting enzyme 2 (ACE-2) (12, 13).

The evolution of overall antibody responses in convalescent individuals is being extensively analysed, with studies showing that Ab titers and neutralization activity against Spike start decreasing during the first weeks after resolution of infection (5, 6, 14-18). Importantly, in addition to neutralizing viral particles, the antiviral activities of SARS-CoV-2-specific antibodies can expand to involve Fc-effector functions, including antibody-dependent cellular cytotoxicity (ADCC) (19, 20). Recent research has highlighted the importance of humoral development and the ability of antibodies to carry out Fc-effector functions in decreasing mortality of patients exhibiting severe disease symptoms (21).

In this study, we dissect multiple aspects of humoral immunity, including Fc-effector functions and antigen-specific B cells, longitudinally for up to 8 months post symptom onset (PSO) in 32 convalescent individuals. Our findings aid in the understanding of durability of COVID-19 immunity, which is important in the context of secondary infections, vaccine efficacy and herd immunity.

## Results

### SARS-CoV-2 RBD-specific and Spike-specific antibody levels in convalescent plasma decrease up to 8 months post-symptom onset

To monitor the evolution of antibody responses longitudinally, we analyzed serological samples from 32 convalescent individuals (Table 1) along with 10 pre-pandemic samples from uninfected individuals as experimental controls. The average age of the donors was 47 years old (range: 20-65 years), and samples were from 17 males and 15 females. Convalescent patients were sampled at four longitudinal time points between 16 and 233 days PSO: 6 weeks (16-95 days; median: 43 days), 11 weeks (48-127 days; median: 77 days), 21 weeks (116-171 days; median: 145 days), and 31 weeks (201-233 days; median: 218 days). Participants were tested positive for SARS-CoV-2 infection by reverse transcription PCR (RT-PCR) on nasopharyngeal swab specimens. Convalescent participants were enrolled following two negative RT-PCR tests and a complete resolution of symptoms for at least 14 days before blood sampling.

**Table 1.** Longitudinal SARS-CoV-2 convalescent cohort

| Group | n | Days after onset of symptoms<br>(median; day range) | Age<br>(median; age range) | Sex |  |
| --- | --- | --- | --- | --- | --- |
|  |  |  |  | Male (n) | Female (n) |
| 6 weeks | 32 | 43<br>(16-95) | 47<br>(20-65) | 17 | 15 |
| 11 weeks | 28 | 77<br>(48-127) | 47<br>(20-65) | 16 | 12 |
| 21 weeks | 28 | 145<br>(116-171) | 48<br>(20-65) | 16 | 12 |
| 31 weeks | 13 | 218<br>(201-233) | 46<br>(20-65) | 9 | 4 |

We began by evaluating the presence of RBD-specific IgG, IgM, and IgA antibodies by using a previously published enzyme-linked immunosorbent assay (ELISA) against the SARS-CoV-2 RBD antigen (16). In agreement with recent reports showing the waning of antibody levels in longitudinal convalescent plasma over time (14-17), we observed that total RBD-specific immunoglobulin (Ig) levels, comprising of IgG, IgM, and IgA, gradually decreased between 6 and 31 weeks after the onset of symptoms (Figure 1A). However, the percentage of convalescent individuals presenting detectable RBD-specific Ig levels remained stable, with a consistent seropositivity rate above 90% throughout the sampling time frame. Notably, 100% of the donors still had detectable IgG at the last time point, while IgM and IgA diminished more rapidly, with 85% and 69% of the donors having undetectable IgM and IgA levels, respectively, 31 weeks PSO (Figure 1B-D; Figure S1A). Since the RBD ELISA is limited to detect antibodies targeting only one domain of the Spike, we developed a high-throughput cell-based ELISA methodology to screen for antibodies recognizing the native full-length S protein on the cell surface. HOS cells stably expressing the SARS-CoV-2 S glycoproteins were incubated with plasma samples, followed by the addition of secondary antibodies recognizing IgG, IgM, and/or IgA. We observed that 100% of the donors still had detectable S-specific total Ig and IgG in their plasma at 31 weeks PSO whereas, only 38% and 54% of the plasma samples tested positive for the presence of Spike-specific IgM and IgA, respectively (Figure 1 E-H; Figure S1B). We confirmed this observation using a recently characterized flow-cytometry based assay (22) determining antibody binding to the full-length S protein on the surface of 293T cells (Figure S2A). The data obtained with both the cell-based ELISA and flow-cytometry techniques correlated significantly (r = 0.8120; p<0.0001) (Figure S2B).

**Figure 1.**
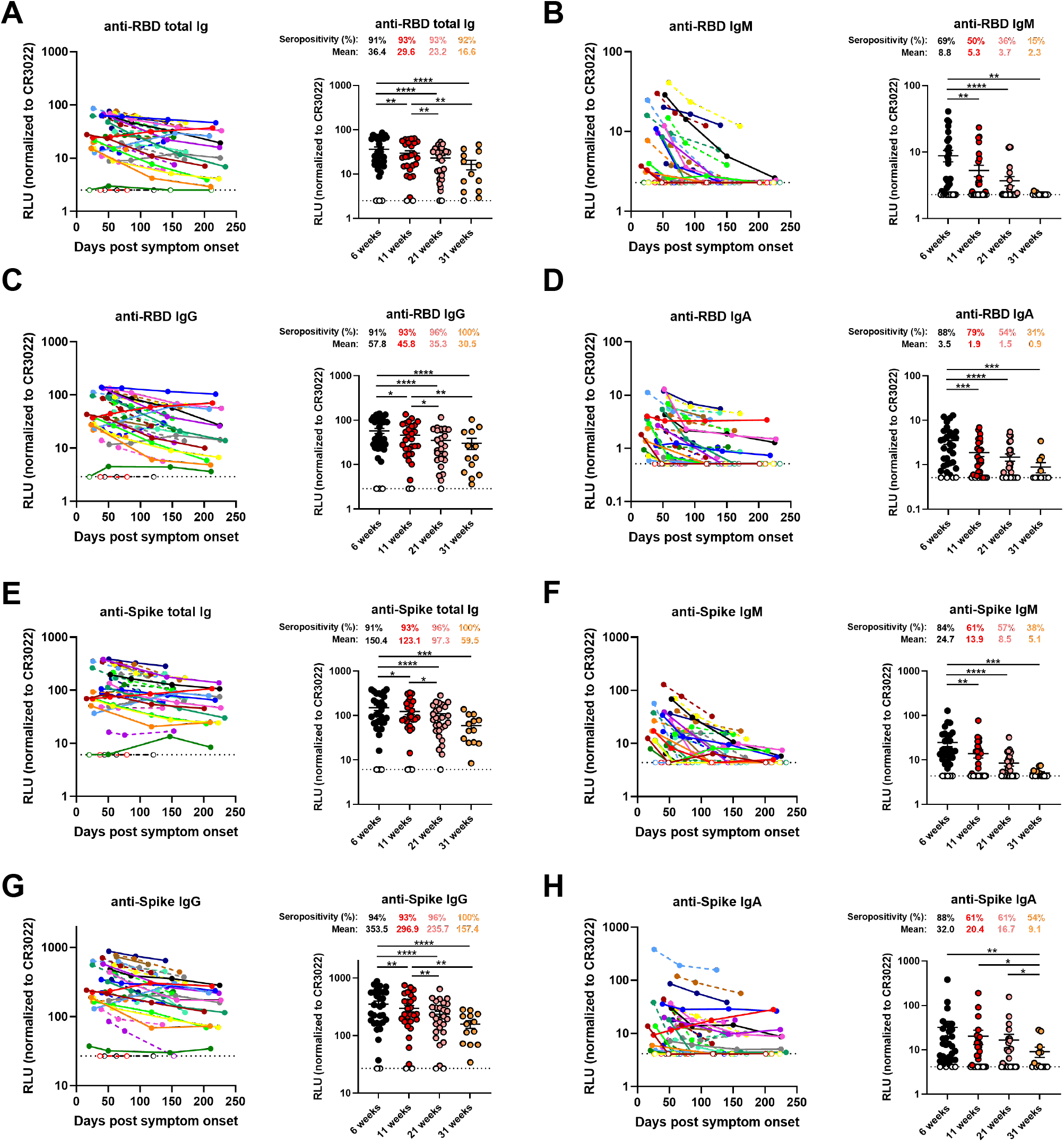
Decline of RBD- and Spike-specific antibodies in longitudinal convalescent plasma. (A-D) Indirect ELISA was performed using recombinant SARS-CoV-2 RBD protein and incubation with COVID-19+ plasma samples recovered between 6 and 31 weeks post-symptom onset. Anti-RBD antibody binding was detected using HRP-conjugated (A) anti-human IgM+IgG+IgA (B) anti-human IgM, (C) anti-human IgG, or (D) anti-human IgA. Relative light unit (RLU) values obtained with BSA (negative control) were subtracted and further normalized to the signal obtained with the anti-RBD CR3022 mAb present in each plate. (E-H) Cell-based ELISA was performed using HOS cells expressing full-length SARS-CoV-2 Spike and incubation with COVID-19+ plasma samples recovered between 6 and 31 weeks post-symptom onset. Anti-Spike antibody binding was detected using HRP-conjugated (E) anti-human IgM+IgG+IgA (F) anti-human IgM, (G) anti-human IgG, or (H) anti-human IgA. RLU values obtained with parental HOS (negative control) were subtracted and further normalized to the signal obtained with the CR3022 mAb present in each plate. (Left panels) Each curve represents the normalized RLUs obtained with the plasma of one donor at every donation as a function of the days after symptom onset. (Right panels) Plasma samples were grouped in different timepoints post-symptom onset (6, 11, 21 and 31 weeks). Undetectable measures are represented as white symbols, and limits of detection are plotted. Error bars indicate means ± SEM. Statistical significance was tested using repeated measures one-way ANOVA with a Holm-Sidak post-test (* P < 0.05; ** P < 0.01; *** P < 0.001; **** P < 0.0001).

### Neutralizing and Fc-effector activities of antibodies present in convalescent plasma decrease at different rates over time

Recent studies have shown the importance of neutralizing antibodies in reducing viral load and preventing infection in animal models (10, 23, 24). Neutralizing monoclonal antibody cocktails also reduced viral load in COVID-19 patients (11). Neutralizing activity is often considered as a determining factor in convalescent plasma therapy, although its relative importance compared to Fc-effector activity is still unknown (25-27). Thus, we measured the capacity of convalescent plasma to neutralize pseudoviral particles carrying the SARS-CoV-2 Spike protein over time. Neutralizing antibody titers (ID_50_) were detected in 63% of the donors at 6 weeks PSO, while none of the uninfected controls had detectable neutralizing activity. Titers declined from 155.6 at 6 weeks to 60.0 at 31 weeks PSO, respectively, with 77% of donors having undetectable neutralization activity in their plasma at the last time point (Figure 2A). Since the depletion of IgM from plasma has been associated with loss of viral neutralization capacity (28, 29) and IgA has also been shown to dominate the early neutralizing response (30, 31), the sharp decline in neutralization activity seen in this study is corroborated by the striking decrease in anti-Spike and anti-RBD IgM and IgA levels (Figure 1B, D, F, H; Figure S1A, B).

**Figure 2.**
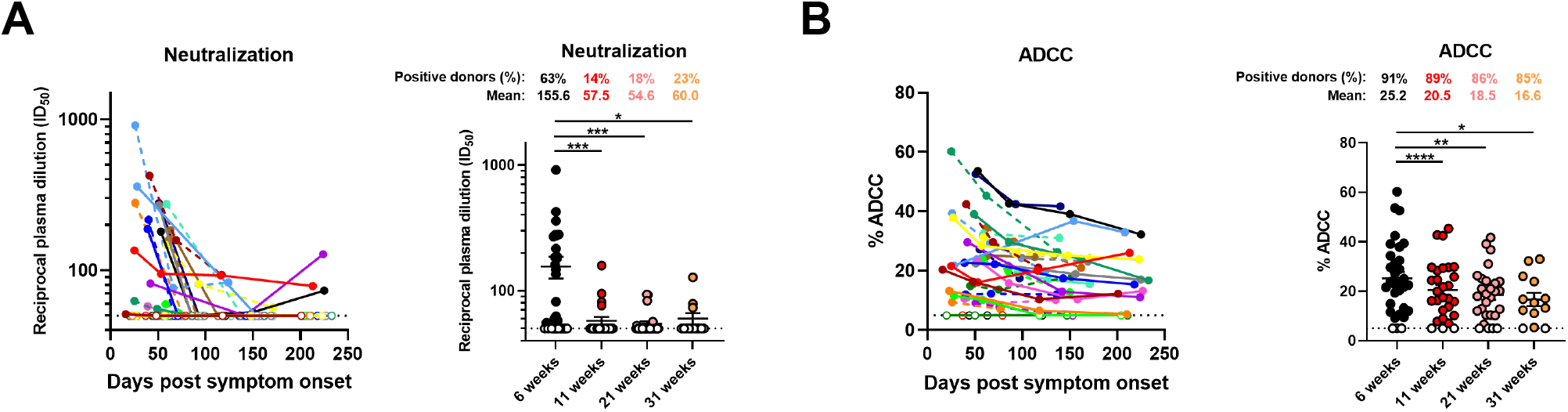
Neutralization and Fc-effector function activities in convalescent plasma decrease over time. (A) Pseudoviral particles coding for the luciferase reporter gene and bearing the SARS-CoV-2 S glycoproteins were used to infect 293T-ACE2 cells. Neutralizing activity was measured by incubating pseudoviruses with serial dilutions of COVID-19+ plasma samples recovered between 6 and 31 weeks post-symptom onset at 37°C for 1 h prior to infection of 293T-ACE2 cells. Neutralization half maximal inhibitory serum dilution (ID_50_) values were determined using a normalized non-linear regression using GraphPad Prism software. (B) CEM.NKr parental cells were mixed at a 1:1 ratio with CEM.NKr-Spike cells and were used as target cells. PBMCs from uninfected donors were used as effector cells in a FACS-based ADCC assay. The graphs shown represent the percentages of ADCC obtained in the presence of COVID-19+ plasma samples recovered between 6 and 31 weeks post-symptom onset. (Left panels) Each curve represents (A) the neutralization ID_50_ or (B) the percentages of ADCC obtained with the plasma of one donor at every donation as a function of the days after symptom onset. (Right panels) Plasma samples were grouped in different timepoints post-symptom onset (6, 11, 21 and 31 weeks). Undetectable measures are represented as white symbols, and limits of detection are plotted. Error bars indicate means ± SEM. Statistical significance was tested using repeated measures one-way ANOVA with a Holm-Sidak post-test (* P < 0.05; ** P < 0.01; *** P < 0.001; **** P < 0.0001).

Fc-mediated effector functions of antibodies can contribute to the efficacy of immune response against SARS-CoV-2. Recent studies have examined Fc-mediated effector functions of antibodies elicited upon SARS-CoV-2 infection (19, 32). Both the presence of IgG and their Fc-mediated effector activities have been linked to reduced severity of disease (21). Herein, we assessed the ability of plasma from convalescent donors to trigger ADCC responses over time. We developed a new ADCC assay using a human T-lymphoid cell line resistant to NK cell-mediated cell lysis (CEM.NKr) and stably expressing the full-length S protein on the cell surface as target cells. PBMCs from healthy individuals were used as effector cells. ADCC activity was measured by the loss of Spike-expressing GFP+ target cells (Figure S3). We observed that ADCC activity of convalescent plasma decreased gradually between 6 weeks and 31 weeks PSO (Figure 2B). However, this decline was modest when compared to the decrease in neutralization activity of plasma (Figure S1C) and 85% of the donors’ plasma still elicited substantial ADCC activity at the latest study time point. The presence of Fc-mediated antibody effector functions up to 8 months PSO is corroborated with the presence of significant IgG levels.

### RBD-specific memory B cells develop and remain stable up to 8 months post-symptom onset

Recent studies on convalescent COVID-19 patients have indicated persistent antigen-specific memory B cell responses despite waning antibody levels (5, 6). To monitor the circulating B cell compartment in our cohort of convalescent individuals, antigen-specific B cells were characterized by flow cytometry (identified as CD19+ CD20+). To identify distinct RBD-specific B cells, we used double discrimination with two recombinant RBD protein preparations labelled with fluorochromes Alexa-Fluor 594 and Alexa-Fluor 488, respectively. Detection of this double positive population was specific since it was not detected in PBMCs from uninfected individuals (Figure S4A, B). RBD-specific B cells were detected 6 weeks PSO with a modest increase in mean frequency up to 31 weeks PSO (0.038% to 0.051%) (Figure 3B). Total RBD-specific B cells were evaluated to distinguish surface Ig isotypes and we observed IgG+, IgM+ and IgA+ cells in 100%, 92% and 83% of donors at 6 weeks PSO, respectively. Strikingly, the frequency of RBD-specific IgM+ B cells decreased significantly, with 46% of the donors having undetectable IgM+ B cells 31 weeks PSO (Figure 3C; Figure S1D). Conversely, the frequency of RBD-specific IgG+ B cells significantly increased between 6 and 21 weeks PSO and remained stable up to 8 months PSO (Figure 3D); the detection of RBD-specific IgA+ B cells also persisted in 77% of the donors tested (Figure 3E; Figure S1D). Furthermore, the total RBD-specific B cells were evaluated to distinguish memory and naïve B cells (identified by using CD27 and CD21 markers; Figure S4C, D). Importantly, total RBD-specific memory B cells were detected in 100% of the donors and the mean frequency remained stable between 6 and 31 weeks PSO (0.020% to 0.026%), while RBD-specific naïve B cells were observed in lower proportions and modestly decreased over time (Figure 3F, G). Interestingly, the proportion of donors positive for RBD-specific naïve B cells was reduced by half between the first and last time point as also observed for RBD-specific IgM+ B cells which is consistent with the decline in IgM levels observed between these groups. IgG+ RBD-specific memory B cells were detected in 100% of the donors and the frequency of this population modestly increased up to 31 weeks PSO (Figure 3H). Meanwhile, the frequencies of IgA+ RBD-specific memory B cells were low but stable over the 8-month period (Figure 3G, I).

**Figure 3.**
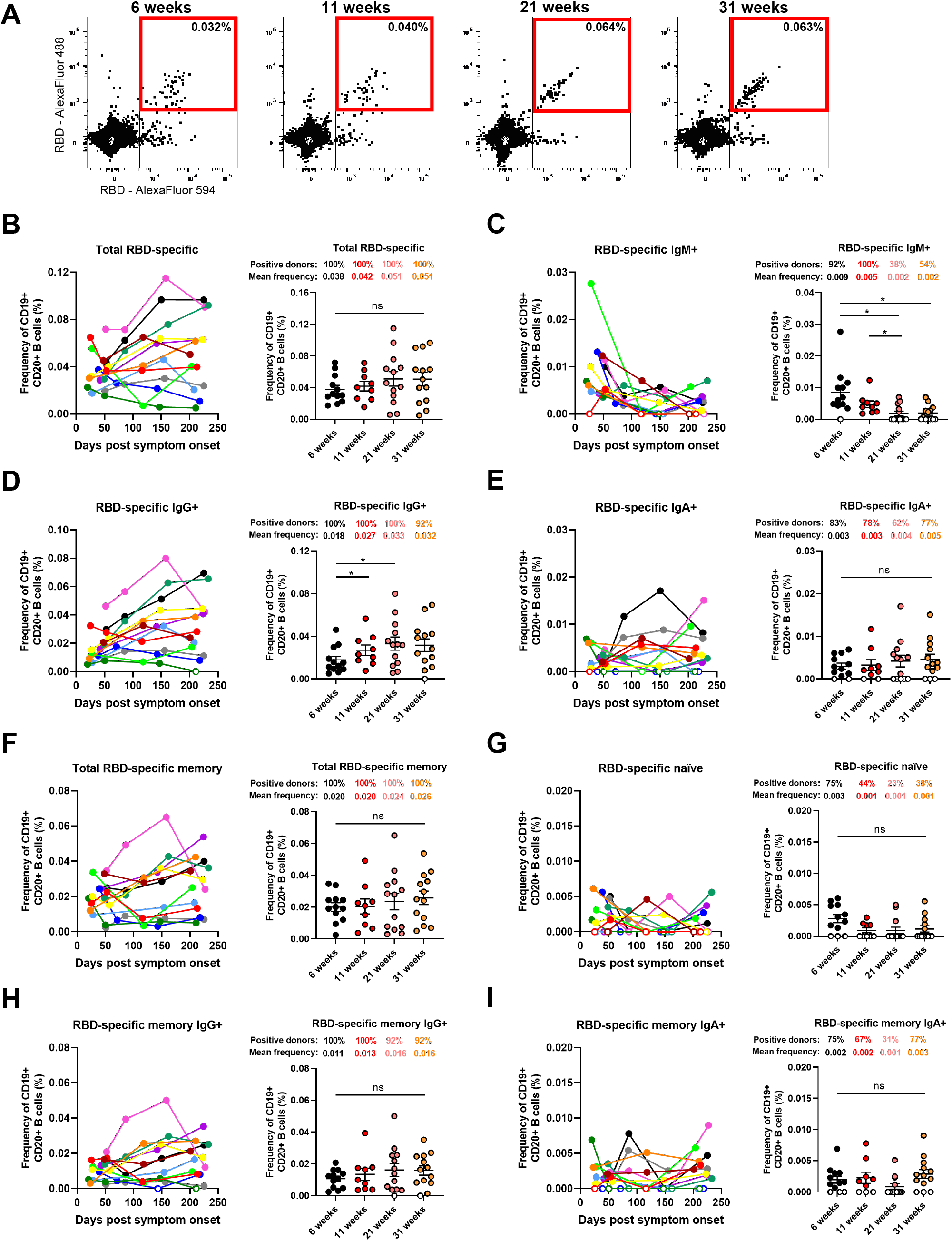
RBD-specific memory B cells develop and persist up to 8 months post-symptom onset. (A) Flow cytometry plots of staining with fluorescent SARS-CoV-2 RBD probes on CD19+ CD20+ HLA-DR+ B cells. Samples from one representative convalescent donor are shown for 4 different timepoints post-symptom onset (6, 11, 21 and 31 weeks). All percentages shown represent the frequency of RBD-specific B cells on the total CD19+ CD20+ B cell population. (B-G) Characterization of RBD-specific B cells was performed on longitudinal PBMC samples obtained from COVID-19+ convalescent individuals between 6 and 31 weeks post-symptom onset. (B) Total RBD-specific B cells were segregated by subsets based on cell surface expression of (C) IgM, (D) IgG or (E) IgA BCR isotypes. Frequency of (F) total memory, (G) naïve, (H) IgG+ memory and (I) IgA+ memory B cells were determined based on CD21 and CD27 expression. (Left panels) Each curve represents the frequency of a B cell subset on the total B cell population obtained with PBMCs from one donor at every donation as a function of the days after symptom onset. (Right panels) PBMC samples were grouped in different timepoints post-symptom onset (6, 11, 21 and 31 weeks). Error bars indicate means ± SEM. Statistical significance was tested using repeated measures one-way ANOVA with a Holm-Sidak post-test (* P < 0.05; ns, nonsignificant)

### Evaluation of the relationship between different aspects of humoral immunity reveals the importance of IgG responses

To examine interrelations between serological, immunological, and functional parameters assessed in this study, we performed comprehensive sets of correlation analyses. Firstly, at each time point, the strongest immune response clusters were comprised of total Ig and IgG binding levels and thus, increased ADCC activity (Figure 4A, left panel). In contrast, weaker immune response clusters in plasma were formed by IgA and IgM levels and subsequently, neutralization capacities. Overall, the responses decreased over the course of the 8-month period (Figure 4B). The inverse was observed with antigen-specific B cell frequencies (Figure 4B). Apart from naïve B cells and the IgM fraction thereof, total RBD-specific B cells increased over the course of 8 months. At each time point, total IgG+ B cells and memory B cells (IgA+ and IgG+) constituted the most prominent responses (Figure 4A, right panel).

**Figure 4.**
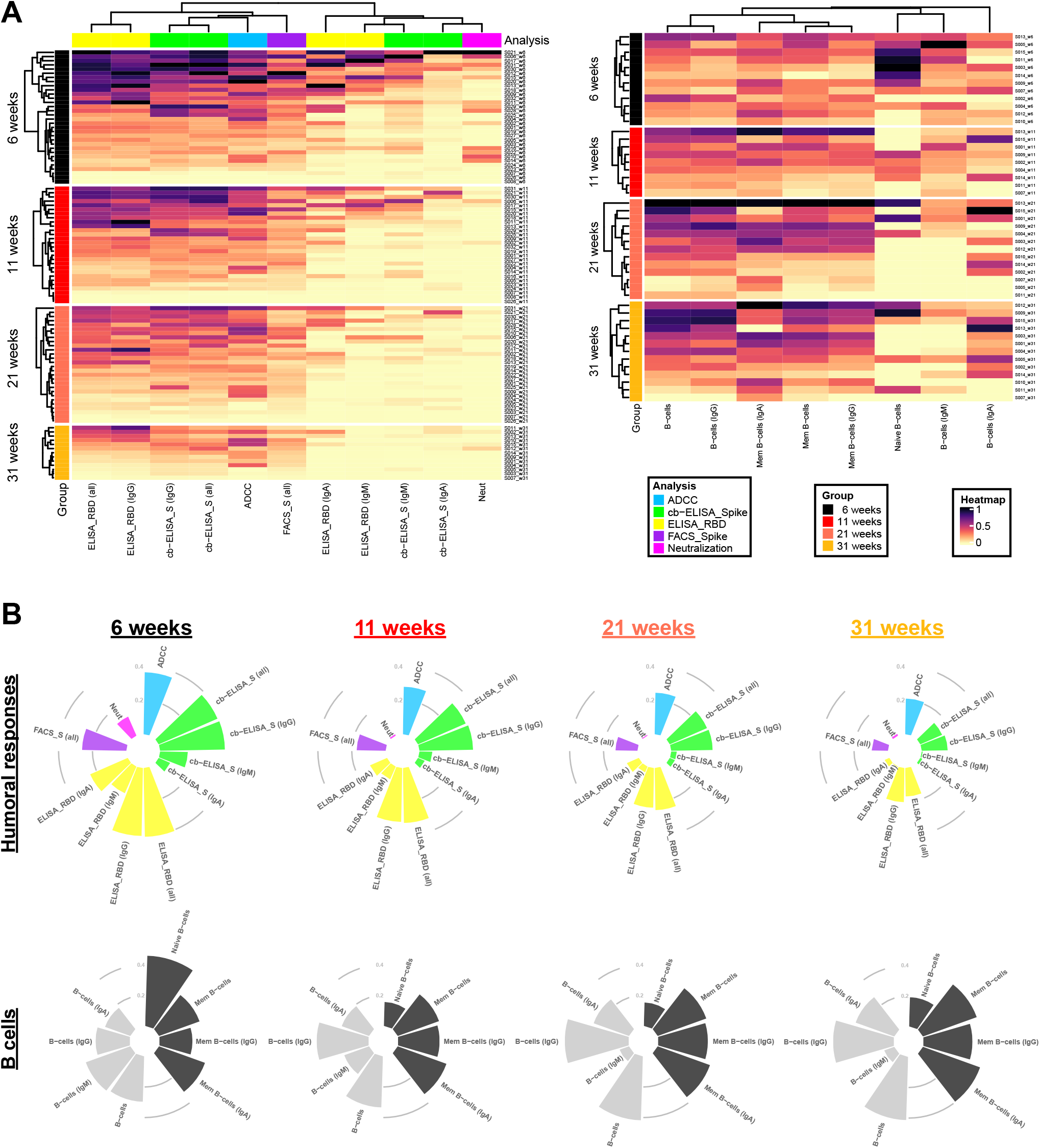
Differential longitudinal patterns of B cell levels and humoral immune responses. (A) (Left panel) Heatmap of humoral immune responses normalized per parameter. Columns represent immune response parameters clustered based on similarity and grouped according to the provided color code. Rows represent IDs grouped according to study time point. IDs are clustered according to their immune response profiles within each time point. (Right panel) Heatmap of B cell levels with similar display as in (left panel). (B) Circular bar plots represent averaged values of parameters at each time point.

Additionally, at the earliest time point convalescent plasma was collected after symptom onset (6 weeks PSO), we observed a vast network of strong positive correlations among B cell populations, antibody levels, and antibody-mediated functional parameters (Figure 5B). Intriguingly, this was followed by a striking disconnect of associations between B cell frequencies and serological parameters at 11, 21, and 31 weeks PSO (Figure 5C-E). This is caused by the concomitant diminution of antibody levels, yet stabilization of antigen-specific B cells observed at these time points, suggestive of decreased antibody production by B cells after resolution of infection or the gradual replacement of Ig-secreting short-lived plasma cell by memory B cells. When considering the collective datasets, prominent features were two subsequently dividing positive correlation clusters, the first among total and IgG+ RBD-specific B cells and the second among ADCC with total Ig and IgG binding responses. Furthermore, days PSO inversely correlated with neutralization as well as IgM and IgA-specific responses (Figure 5A and S5).

**Figure 5.**
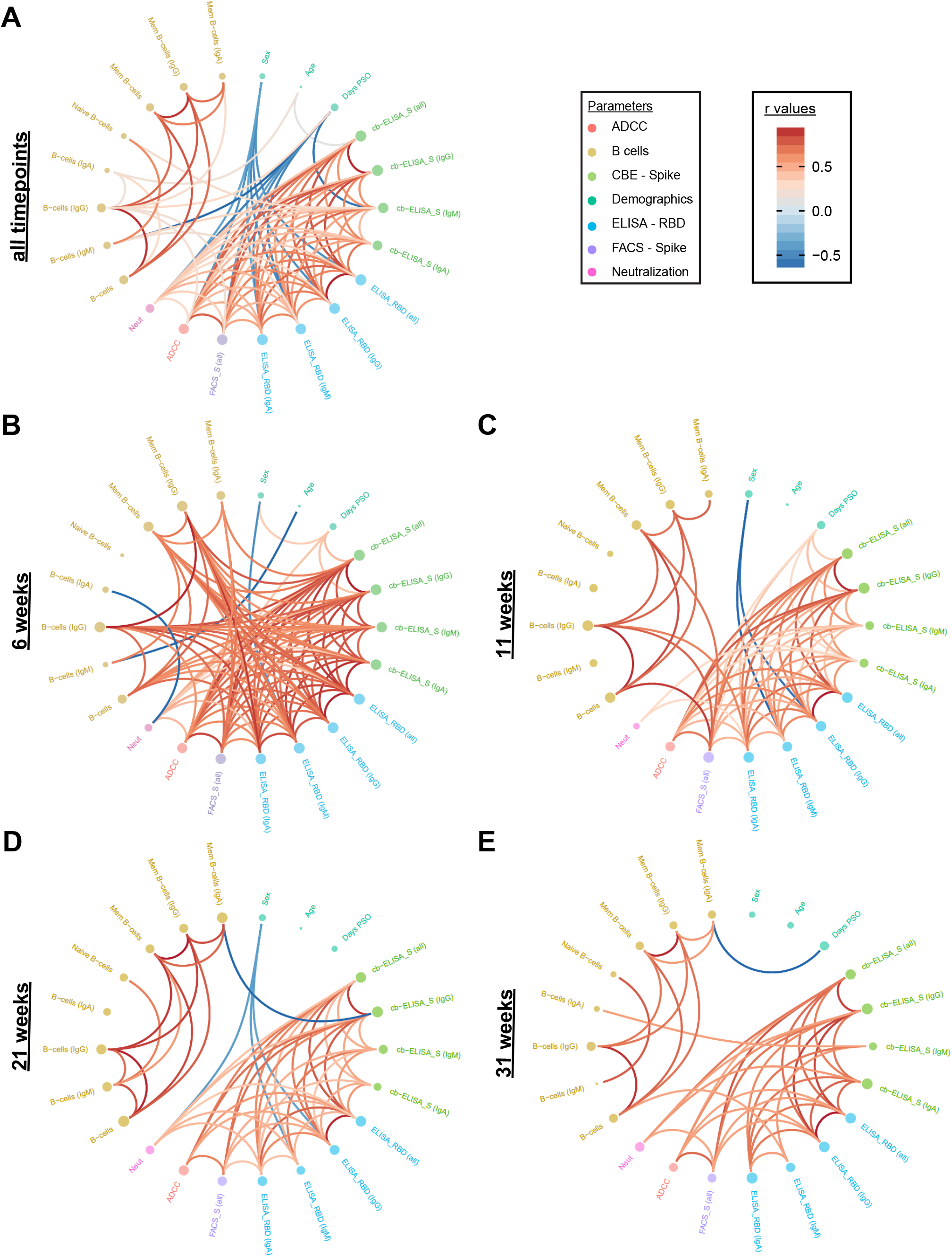
Longitudinal plasticity and separation of B cell and humoral correlation clusters. Edge bundling correlation plots where red and blue edges represent positive and negative correlations between connected parameters, respectively. Only significant correlations (p < 0.05) are displayed. Nodes are color-coded based on the grouping of parameters according to the legend at the bottom. Node size corresponds to the degree of relatedness of correlations. Edge bundling plots are shown for correlation analyses using all data points (A) and datasets of individual time points, i.e., at 6 weeks (B), 11 weeks (C), 21 weeks (D), and 31 weeks (E).

## Discussion

A better understanding of the type and longevity of immune responses following viral infections are critical to reveal immune mechanisms involved in protection from re-infection and protection by vaccination. Data regarding these important issues continues to be gathered for SARS-CoV-2. In this study, we have contributed to the current understanding by reporting on the evolution of the overall humoral immune responses in 101 blood samples obtained from 32 COVID-19 convalescent patients between 16 and 233 days PSO. Overall, we observed that IgM levels and the neutralizing capacity of plasma decreases rapidly, whereas IgG and Fc-effector activity are more sustained (Figure 6). The antibody kinetics we observed herein are typical of those seen for other human coronavirus infections, with antibodies peaking 2-4 weeks PSO followed by a contraction phase (33). Furthermore, we show that COVID-19 patients generate RBD-specific memory B cells and IgG+ B memory cells that persist for over 8 months. Similarly, recent studies on the durability of SARS-CoV-2 immune responses have shown that although antibody levels decrease, Spike-specific IgG+ memory B cell responses are generated and maintained (1, 5, 6). Additionally, studies demonstrated enhanced cellular immunity that protects non-human primates from SARS-CoV-2 reinfection in the context of waning neutralizing antibodies (8). Thus, the decline of antibody levels does not negate the protective potential because of the importance of cellular responses against SARS-CoV-2 infection. This understanding is corroborated with a recent risk assessment carried out in a cohort of 43,000 convalescent individuals demonstrating that immunity elicited upon natural SARS-CoV-2 infection protects against reinfection with an efficacy of >90% for at least 7 months (34).

**Figure 6.**
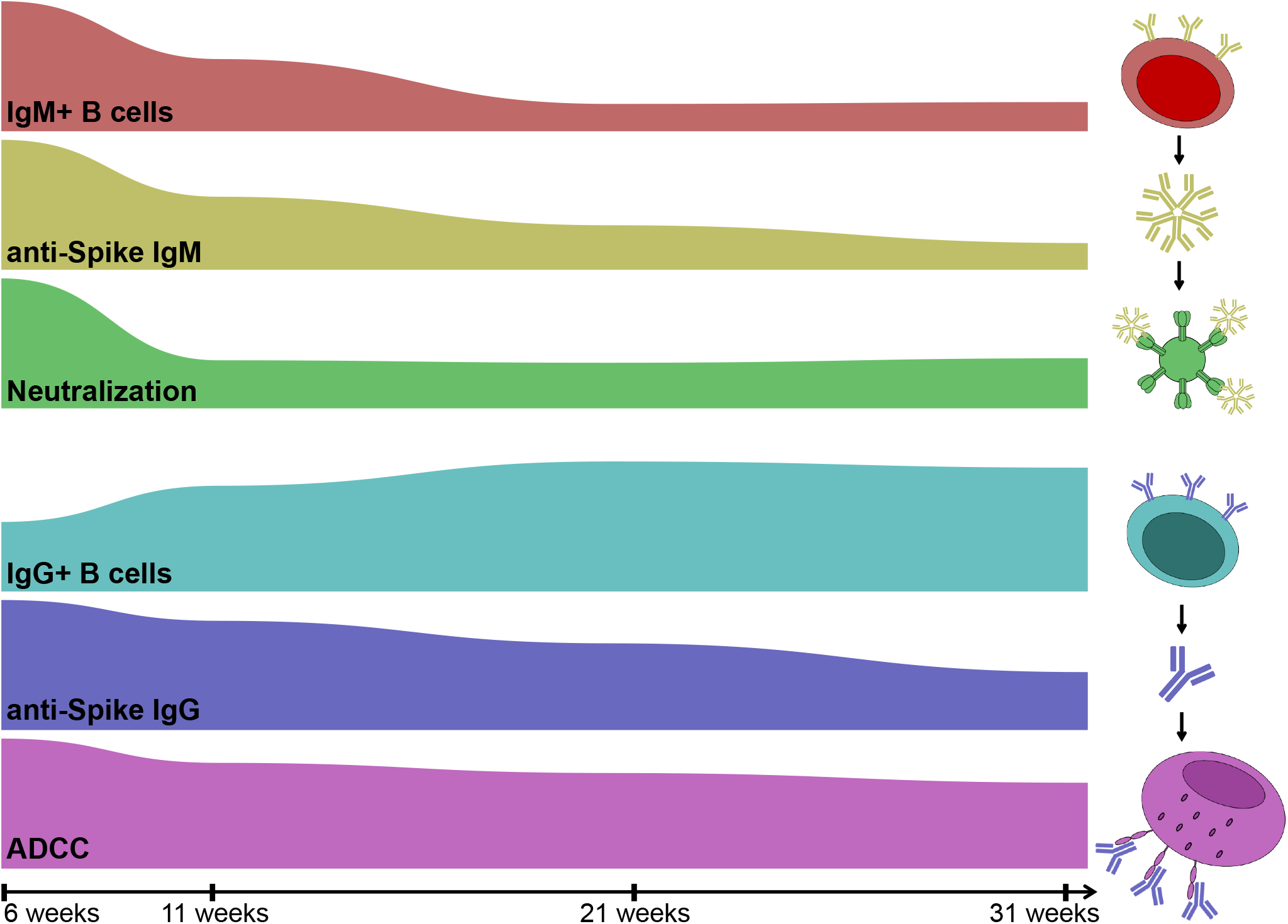
Evolution of humoral immune responses and B cell levels in SARS-CoV-2 convalescent individuals over time. Area plots showing time series of selected, humoral immune responses and B cell levels by interpolation of normalized, averaged values per parameter and time point. The timeline is shown at the bottom with ticks indicating study time points.

Fc-mediated effector activity of antibodies has been recently shown to correlate with reduced disease severity and mortality (21). Antibodies capable of mediating Fc-dependant functions, such as antibody-dependent phagocytosis and ADCC, have been isolated from convalescent donors (35). Importantly, Fc-mediated effector activity was shown to protect from SARS-CoV-2 infection in adapted mice and hamster models (9, 19, 32). Thus, our observations of the persistence of ADCC responses in convalescent plasma up to 8 months PSO suggest that Fc-mediated effector functions could play a vital role in protection from reinfection. Nevertheless, the heterogeneity of immune responses between individuals is and will be an important factor when evaluating the efficacy of immune responses upon re-exposure to SARS-CoV-2.

A recent study comparing humoral immunity generated by mRNA vaccinees (mRNA-1273 or BNT162b2) and individuals recovered from natural infection observed similarities in antibody binding titers and plasma neutralization capacity (36). Furthermore, this study also observed similar frequencies of RBD-specific memory B cells between vaccinees and infected individuals. Therefore, our results on the persistence of RBD-specific memory B cells up to 8 months after natural infection is reassuring with regards to long-term vaccine efficacy.

## Material and Methods

### Ethics Statement

All work was conducted in accordance with the Declaration of Helsinki in terms of informed consent and approval by an appropriate institutional board. Convalescent plasmas were obtained from donors who consented to participate in this research project at CHUM (19.381). The donors met all donor eligibility criteria: previous confirmed COVID-19 infection and complete resolution of symptoms for at least 14 days.

### Plasma and antibodies

Plasma from SARS-CoV-2-infected and pre-pandemic uninfected donors were collected, heat-inactivated for 1h at 56°C and stored at -80°C until ready to use in subsequent experiments. Plasma from uninfected donors were used as negative controls and used to calculate the seropositivity threshold in our ELISA, cell-based ELISA, and flow cytometry assays. The RBD-specific monoclonal antibody CR3022 was used as a positive control in our ELISAs, cell-based ELISAs, and flow cytometry assays and was previously described (14, 16, 22). Horseradish peroxidase (HRP)-conjugated antibodies able to detect all Ig isotypes (anti-human IgM+IgG+IgA; Jackson ImmunoResearch Laboratories, Inc.) or specific for the Fc region of human IgG (Invitrogen), the Fc region of human IgM (Jackson ImmunoResearch Laboratories, inc.) or the Fc region of human IgA (Jackson ImmunoResearch Laboratories, inc) were used as secondary antibodies to detect antibody binding in ELISA and cell-based ELISA experiments. Alexa Fluor-647-conjugated goat anti-human Abs able to detect all Ig isotypes (anti-human IgM+IgG+IgA; Jackson ImmunoResearch Laboratories, Inc.) were used as secondary antibodies to detect plasma binding in flow cytometry experiments.

### Cell lines

293T human embryonic kidney cells (obtained from ATCC) were maintained at 37°C under 5% CO_2_ in Dulbecco’s modified Eagle’s medium (DMEM) (Wisent) containing 5% fetal bovine serum (VWR) and 100 μg/ml penicillin-streptomycin (Wisent). 293T-ACE2 and 293T-SARS-CoV-2 Spike cell lines were previously reported (16). For the generation of HOS and CEM.NKr CCR5+ cells stably expressing the SARS-CoV-2 Spike glycoproteins, transgenic lentiviruses were produced in 293T using a third-generation lentiviral vector system. Briefly, 293T cells were co-transfected with two packaging plasmids (pLP1 and pLP2), an envelope plasmid (pSVCMV-IN-VSV-G) and a lentiviral transfer plasmid coding for a GFP-tagged SARS-CoV-2 Spike (pLV-SARS-CoV-2 S C-GFPSpark tag) (Sino Biological). Supernatant containing lentiviral particles was used to transduce HOS and CEM.NKr CCR5+ cells in presence of 5μg/mL polybrene. The HOS and CEM.NKr CCR5+ cells stably expressing SARS-CoV-2 Spike (GFP+) were sorted by flow cytometry.

### Protein expression and purification

FreeStyle 293F cells (Invitrogen) were grown in FreeStyle 293F medium (Invitrogen) to a density of 1 × 10^6^ cells/mL at 37°C with 8% CO_2_ with regular agitation (150 rpm). Cells were transfected with a plasmid coding for SARS-CoV-2 S RBD using ExpiFectamine 293 transfection reagent, as directed by the manufacturer (Invitrogen). One week later, cells were pelleted and discarded. Supernatants were filtered using a 0.22 µm filter (Thermo Fisher Scientific). The recombinant RBD proteins were purified by nickel affinity columns, as directed by the manufacturer (Invitrogen). The RBD preparations were dialyzed against phosphate-buffered saline (PBS) and stored in aliquots at -80°C until further use. To assess purity, recombinant proteins were loaded on SDS-PAGE gels and stained with Coomassie Blue.

### Enzyme-Linked Immunosorbent Assay (ELISA)

The SARS-CoV-2 RBD ELISA assay used was recently described (16). Briefly, recombinant SARS-CoV-2 S RBD proteins (2.5 μg/ml), or bovine serum albumin (BSA) (2.5 μg/ml) as a negative control, were prepared in PBS and adsorbed to plates (MaxiSorp Nunc) overnight at 4°C. Coated wells were subsequently blocked with blocking buffer (Tris-buffered saline [TBS] containing 0.1% Tween20 and 2% BSA) for 1h at room temperature. Wells were then washed four times with washing buffer (Tris-buffered saline [TBS] containing 0.1% Tween20). CR3022 mAb (50ng/ml) or a 1/250 dilution of plasma from SARS-CoV-2-infected or uninfected donors were prepared in a diluted solution of blocking buffer (0.1 % BSA) and incubated with the RBD-coated wells for 90 minutes at room temperature. Plates were washed four times with washing buffer followed by incubation with secondary Abs (diluted in blocking buffer,0.4% BSA) for 1h at room temperature, followed by four washes. HRP enzyme activity was determined after the addition of a 1:1 mix of Western Lightning oxidizing and luminol reagents (Perkin Elmer Life Sciences). Light emission was measured with a LB942 TriStar luminometer (Berthold Technologies). Signal obtained with BSA was subtracted for each plasma and was then normalized to the signal obtained with CR3022 mAb present in each plate. The seropositivity threshold was established using the following formula: mean of all COVID-19 negative plasma + (3 standard deviation of the mean of all COVID-19 negative plasma).

### Cell-Based ELISA

Detection of the trimeric SARS-CoV-2 Spike at the surface of HOS cells was performed by cell-based enzyme-linked immunosorbent assay (ELISA). Briefly, parental HOS cells or HOS-Spike cells were seeded in 384-well plates (2.8×10^4^ cells per well) overnight. Cells were blocked with blocking buffer (washing buffer [1.8 mM CaCl_2_, 1 mM MgCl_2_, 25 mM Tris (pH 7.5), and 140 mM NaCl] supplemented with 10mg/mL non-fat dry milk and 5mM Tris [pH 8.0]) for 30min. CR3022 mAb (1 μg/ml) or plasma from SARS-CoV-2-infected or uninfected donors (at a dilution of 1/250) were prepared in blocking buffer and incubated with the cells for 1h at room temperature. Respective HRP-conjugated secondary antibodies were then incubated with the samples for 45 min at room temperature. For all conditions, cells were washed 6 times with blocking buffer and 6 times with washing buffer. HRP enzyme activity was determined after the addition of a 1:1 mix of Western Lightning oxidizing and luminol reagents (PerkinElmer Life Sciences). Light emission was measured with an LB 942 TriStar luminometer (Berthold Technologies). Signal obtained with parental HOS was subtracted for each plasma and was then normalized to the signal obtained with CR3022 mAb present in each plate. The seropositivity threshold was established using the following formula: mean of all COVID-19 negative plasma + (3 standard deviation of the mean of all COVID-19 negative plasma).

### Cell surface staining and flow cytometry analysis

293T and 293T-Spike cells were mixed at a 1:1 ratio and stained with the anti-RBD CR3022 monoclonal Ab (5 μg/ml) or plasma (1:250 dilution). AlexaFluor-647-conjugated goat anti-human IgM+IgG+IgA Abs (1:800 dilution) were used as secondary antibodies. The percentage of transduced cells (GFP+ cells) was determined by gating the living cell population based on viability dye staining (Aqua Vivid, Invitrogen). Samples were acquired on a LSRII cytometer (BD Biosciences) and data analysis was performed using FlowJo v10.7.1 (Tree Star). The seropositivity threshold was established using the following formula: mean of all COVID-19 negative plasma + (3 standard deviation of the mean of all COVID-19 negative plasma).

### ADCC assay

For evaluation of anti-SARS-CoV-2 antibody-dependent cellular cytotoxicity (ADCC), parental CEM.NKr CCR5+ cells were mixed at a 1:1 ratio with CEM.NKr. Spike cells. These cells were stained for viability (AquaVivid; Thermo Fisher Scientific, Waltham, MA, USA) and cellular dyes (cell proliferation dye eFluor670; Thermo Fisher Scientific) and subsequently used as target cells. Overnight rested PBMCs were stained with another cellular marker (cell proliferation dye eFluor450; Thermo Fisher Scientific) and used as effector cells. Stained target and effector cells were mixed at a ratio of 1:10 in 96-well V-bottom plates. Plasma from COVID+ or COVID-individuals (1/500 dilution) or monoclonal antibody CR3022 (1 µg/mL) were added to the appropriate wells. The plates were subsequently centrifuged for 1 min at 300xg, and incubated at 37°C, 5% CO_2_ for 5 hours before being fixed in a 2% PBS-formaldehyde solution. ADCC activity was calculated using the formula: [(% of GFP+ cells in Targets plus Effectors)-(% of GFP+ cells in Targets plus Effectors plus plasma/antibody)]/(% of GFP+ cells in Targets) x 100 by gating on transduced live target cells. All samples were acquired on an LSRII cytometer (BD Biosciences) and data analysis performed using FlowJo v10.7.1 (Tree Star). The specificity threshold was established using the following formula: mean of all COVID-19 negative plasma + (3 standard deviation of the mean of all COVID-19 negative plasma)

### Virus neutralization assay

293T-ACE2 target cells were infected with single-round luciferase-expressing SARS-CoV-2 pseudoparticles in presence of convalescent plasma. Briefly, 293T cells were transfected by the calcium phosphate method with the lentiviral vector pNL4.3 R-E-Luc (NIH AIDS Reagent Program) and a plasmid encoding for SARSCoV-2 Spike at a ratio of 5:4. Two days post-transfection, cell supernatants were harvested and stored at -80°C until use. 293T-ACE2 target cells were seeded at a density of 1×10^4^ cells/well in 96-well luminometer-compatible tissue culture plates (Perkin Elmer) 24h before infection. Recombinant viruses in a final volume of 100 µL were incubated with the indicated plasma dilutions (1/50; 1/250; 1/1250; 1/6250; 1/31250) for 1h at 37°C and were then added to the target cells followed by incubation for 48h at 37°C; cells were lysed by the addition of 30 µL of passive lysis buffer (Promega) followed by one freeze-thaw cycle. An LB942 TriStar luminometer (Berthold Technologies) was used to measure the luciferase activity of each well after the addition of 100 µL of luciferin buffer (15mM MgSO_4_, 15mM KPO_4_ [pH 7.8], 1mM ATP, and 1mM dithiothreitol) and 50 µL of 1mM d-luciferin potassium salt (Thermo Fisher Scientific). The neutralization half-maximal inhibitory dilution (ID_50_) represents the plasma dilution to inhibit 50% of the infection of 293T-ACE2 cells by SARS-CoV-2 pseudoviruses.

### Detection of antigen-specific B cells

To detect SARS-CoV-2-specific B cells, we conjugated recombinant RBD proteins with Alexa Fluor 488 or Alexa Fluor 594 (Thermo Fisher Scientific) according to the manufacturer’s protocol. Approximately 10 × 10^6^ frozen PBMC from 13 convalescent donors were prepared in Falcon® 5ml-round bottom polystyrene tubes at a final concentration of 14 × 10^6^ cells/mL in RPMI 1640 medium (Gibco by Life Technologies, #11875-093) supplemented with 10% of fetal bovine serum (Seradigm, #1500-500), Penicillin-Streptomycin (Gibco by Life Technologies, #15140122) and HEPES (Gibco by Life Technologies, #15630-080). After a rest of 2h at 37°C and 5% CO_2_, cells were stained using Aquavivid viability marker (Gibco by Life Technologies) in DPBS (Gibco by Life Technologies, #14190-144) at 4°C for 20min. The detection of SARS-CoV-2-antigen specific B cells was done by adding the RBD probes to the following antibody cocktail: IgM BUV737 (Clone UCH-B1, #748928), CD24 BUV805 (Clone ML5, #742010), IgG BV421 (Clone G18-145, #562581), CD3 BV480 (Clone UCHT1, #), CD56 BV480 (Clone NCAM16.2, #566124), CD14 BV480 (Clone NCAM16.2, #746304), CD16 BV480 (Clone 3G8, #566108), CD20 BV711 (Clone 2H7, #563126), CD21 BV786 (Clone B-LY4, #740969), HLA DR BB700 (Clone G46-6, #566480), CD27 APC R700 (Clone M-T271, #565116) all from BD Biosciences; CD19 BV650 (Clone SJ25C1, #363026) from Biolegend and IgA PE (Clone IS11-8E10, #130-113-476) from Miltenyi. Staining was performed at 4°C for 30min and cells were fixed using 2% paraformaldehyde at 4°C for 15min. Stained PBMC samples were acquired on Symphony cytometer (BD Biosciences) and analyzed using FlowJo v10.7.1 (TreeStar). In each experiment, PBMC from unexposed donors (total of n=9) were included to ensure consistent specificity of the assay.

### Statistical analyses

Statistics were analyzed using GraphPad Prism version 9.0.0 (GraphPad, San Diego, CA). Every dataset was tested for statistical normality and this information was used to apply the appropriate (parametric or nonparametric) statistical test. P values < 0.05 were considered significant; significance values are indicated as *p < 0.05, **p < 0.01, ***p < 0.001, ****p < 0.0001. Multiplicity adjustments of p values were performed with the Benjamini-Hochberg method in R and R Studio (37, 38) using the data.table and tidyverse packages.

### Software scripts and visualization

Normalized heatmaps were generated using the complexheatmap, tidyverse, and viridis packages in R and RStudio (37, 38). Normalizations were done per parameter. IDs were grouped and clustered separately according to time point. Correlograms were generated using the corrplot and RColorBrewer packages in program R and RStudio using hierarchical clustering according to the first principal component (FPC). Circular barplots were generated in R and RStudio using the tidyverse package with averaged, normalized data per parameter and time point. Edge bundling graphs were generated in undirected mode in R and RStudio using ggraph, igraph, tidyverse, and RColorBrewer packages. Edges are only shown if p < 0.05, and nodes are sized according to the connecting edges’ r values. Nodes are color-coded according to groups of parameters. Area graphs were generated for the display of normalized time series. The plots were created in RawGraphs using DensityDesign interpolation and vertically un-centered values (39).

## Acknowledgements

The authors are grateful to the convalescent plasma donors who participated in this study. The authors thank the CRCHUM BSL3 and Flow Cytometry Platforms for technical assistance. We thank Dr. Stefan Pöhlmann and Dr. Markus Hoffmann (Georg-August University, Germany) for the plasmid coding for SARS-CoV-2 S glycoproteins and Dr. M. Gordon Joyce (U.S. MHRP) for the monoclonal antibody CR3022. This work was supported by le Ministère de l’Économie et de l’Innovation du Québec, Programme de soutien aux organismes de recherche et d’innovation to A.F., by the Fondation du CHUM and the Fondation du CHU Sainte-Justine. This work was also supported by Canada’s COVID-19 Immunity Task Force (CITF), in collaboration with the Canadian Institutes of Health Research (CIHR) (Grant VR2-173203), a CIHR foundation grant #352417 to A.F., by CIHR COVID-19 Rapid Research Funding to A.F., R.B. and P.B. and by an Exceptional Fund COVID-19 from the Canada Foundation for Innovation (CFI) #41027 to A.F. and D.E.K. A.F. is the recipient of Canada Research Chair on Retroviral Entry no. RCHS0235 950-232424. V.M.L. and P.B. are supported by FRQS Junior 1 salary awards. D.E.K. is a FRQS Merit Research Scholar. R.D. was supported by NIH grant R01 AI122953-05. S.P.A, J.P. and G.B.B. are supported by CIHR fellowships. R.G. is supported by a MITACS Accélération postdoctoral fellowship. The funders had no role in study design, data collection and analysis, decision to publish, or preparation of the manuscript. We declare no competing interests.

---

**Supplementary Figure 1.**
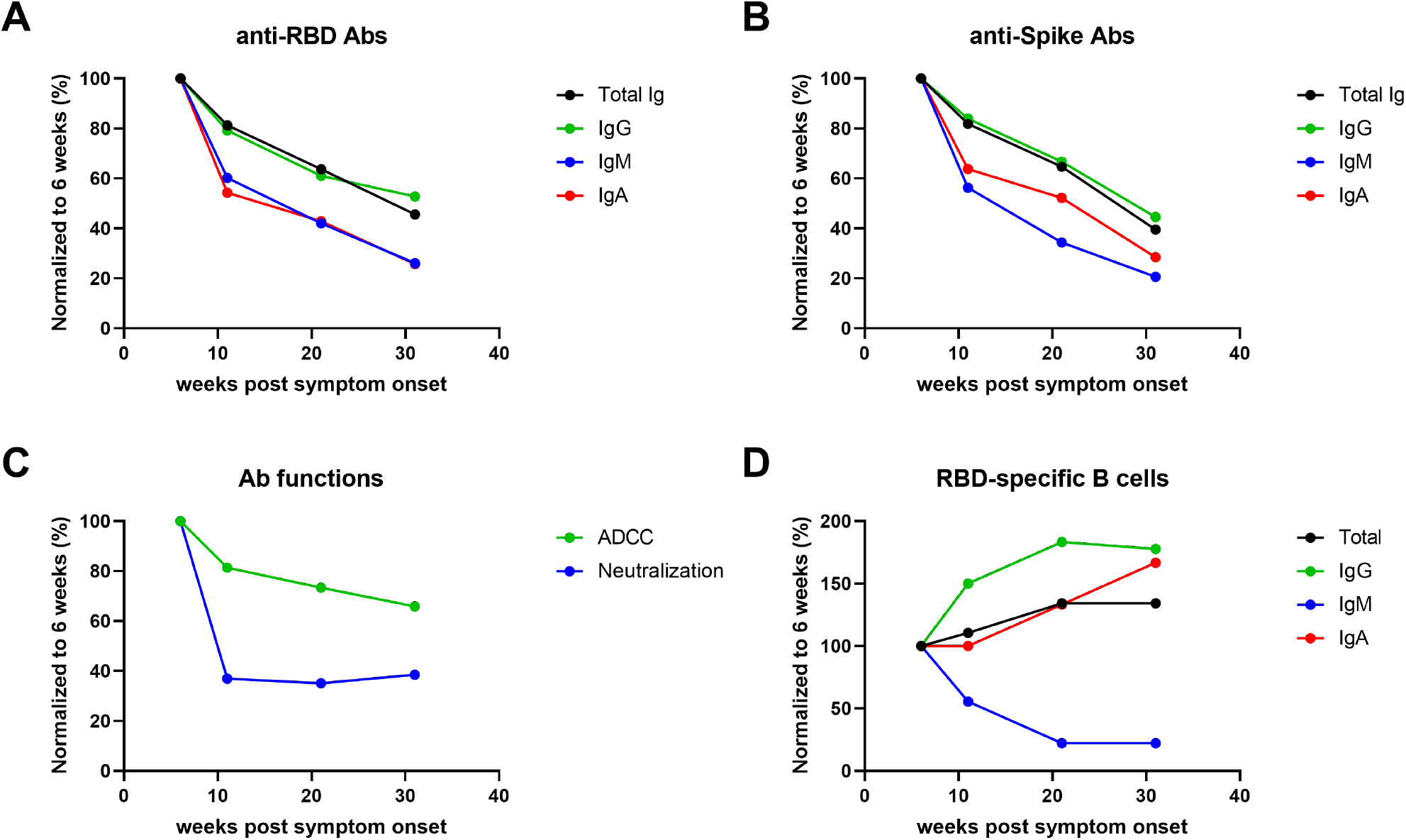
Anti-SARS-CoV-2 IgM and IgA levels decline faster than IgG in the convalescence phase. (A) The graph shown represents the mean values for anti-RBD ELISA (from Figure 1A-D) at different timepoints (6, 11, 21 and 31 weeks) normalized to the 6 weeks timepoint. (B) The graph shown represents the mean values for anti-Spike cell-based ELISA (from Figure 1E-H) at different timepoints (6, 11, 21 and 31 weeks) normalized to the 6 weeks timepoint. (C) The graph shown represents the mean values for neutralization and ADCC responses (from Figure 2) at different timepoints (6, 11, 21 and 31 weeks) normalized to the 6 weeks timepoint. (D) The graph shown represents the mean values for RBD-specific B cell frequencies (from Figure 3B-E) at different timepoints (6, 11, 21 and 31 weeks) normalized to the 6 weeks timepoint.

**Supplementary Figure 2.**
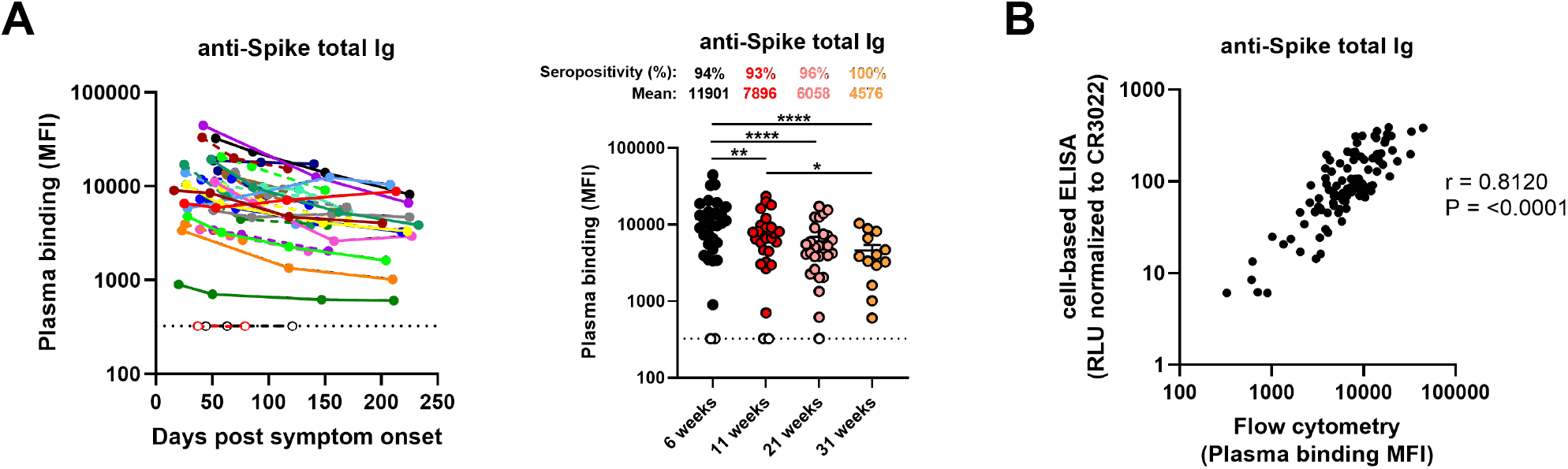
Detection of antibodies against SARS-CoV-2 Spike by flow cytometry correlates with anti-Spike detection by cell-based ELISA. (A) Cell-surface staining of 293T cells stably expressing full-length SARS-CoV-2 Spike using samples from COVID-19+ convalescent donors at different times after symptoms onset (6, 11, 21 and 31 weeks). The graphs shown represent the median fluorescence intensities (MFI) obtained on the GFP+ population. MFIs values obtained with parental 293T (GFP-) were subtracted. (Left panel) Each curve represents the MFIs obtained with the plasma of one donor at every donation as a function of the days after symptom onset. (Right panel) Plasma samples were grouped in different timepoints post-symptom onset (6, 11, 21 and 31 weeks). Undetectable measures are represented as white symbols, and limits of detection are plotted. Error bars indicate means ± SEM. (B) The levels of anti-Spike total Ig quantified by flow cytometry were correlated with the level of anti-Spike total Ig quantified by cell-based ELISA. Statistical significance was tested using (A) a repeated measures one-way ANOVA with a Holm-Sidak post-test or (B) a Spearman correlation rank test (* P < 0.05; ** P < 0.01; **** P < 0.0001).

**Supplementary Figure 3.**
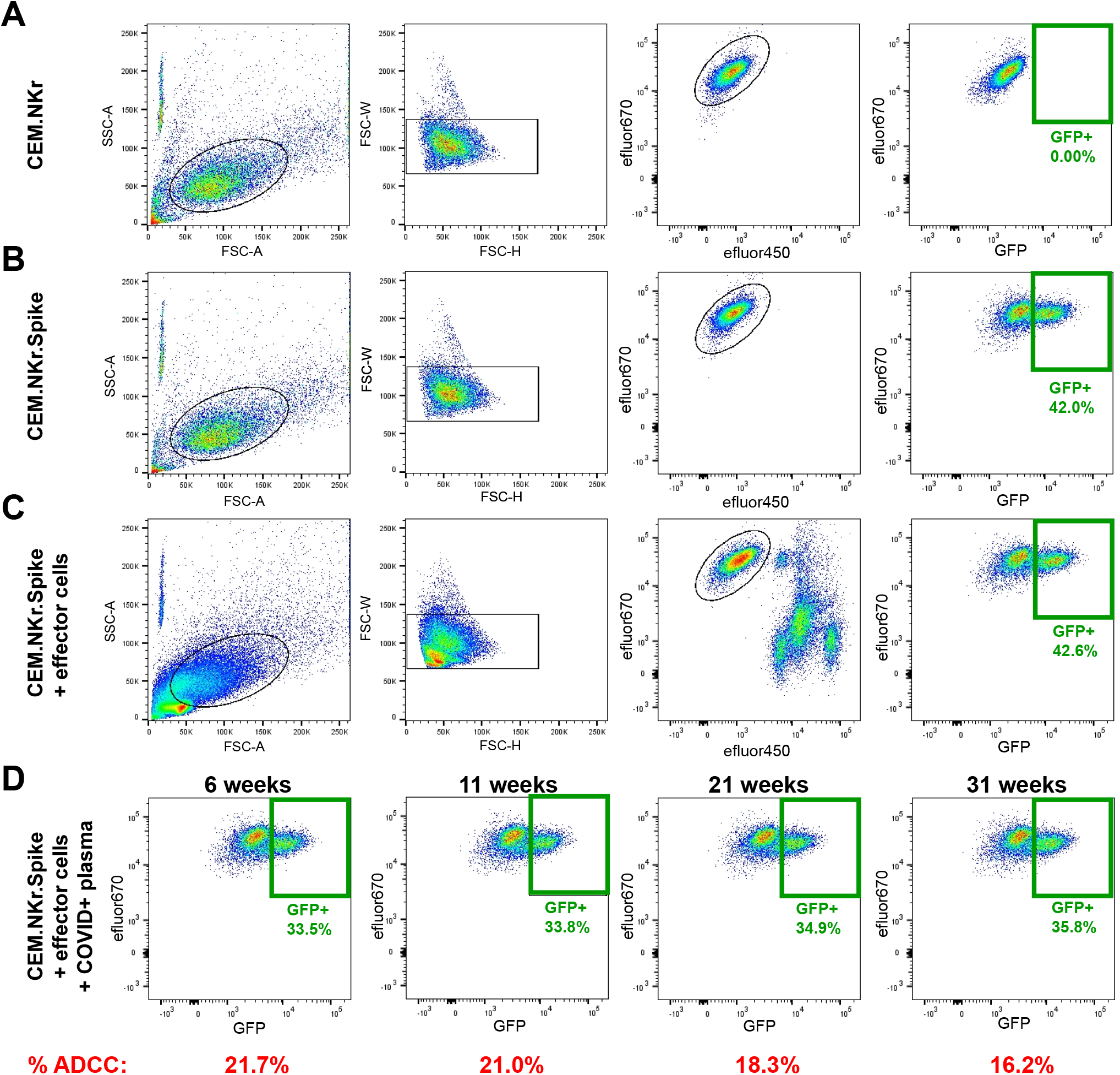
Gating strategy for ADCC measurements. (A) Target cells were identified according to cell morphology by light-scatter parameters (first column) and excluding doublets cells (second column). Cells were then gated on eFluor670+ cells (excluding the effector cells labeled with eFluor450; third column). Finally, the percentage of GFP+ target cells was used to calculate ADCC activity (last column). Examples of gating using (A) parental CEM.NKr or (B) a 1:1 ratio mix of CEM.NKr and CEM.NKr.Spike as target cells in absence or (C) in presence of effector cells. (D) ADCC assay performed in the presence of plasma samples from one representative convalescent donor at 4 different timepoints post-symptom onset (6, 11, 21 and 31 weeks).

**Supplementary Figure 4.**
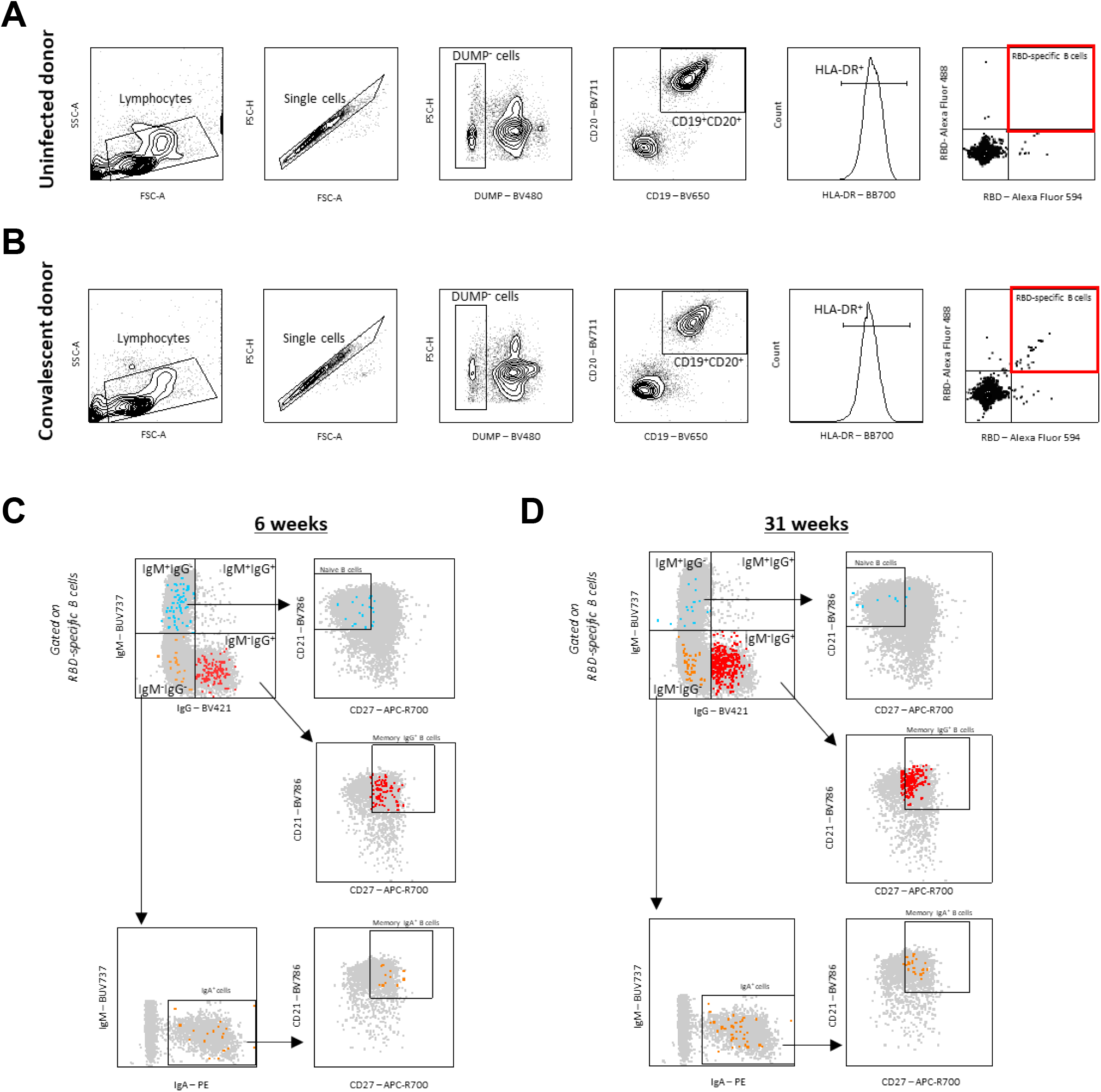
Gating strategy for SARS-CoV-2-specific B cell characterization. (A-B) Representative flow cytometry gates to identify RBD-specific B cells from PBMCs of (A) uninfected and (B) convalescent donor. (C-D) Flow cytometry gates used to differentiate RBD-specific B cell subtypes using isotypic and maturation cell surface markers on samples obtained (C) 6 weeks and (D) 31 weeks post-symptom onset. After identification of isotypic subtypes, RBD-specific naïve and memory B cells were characterized based on surface expression of CD21 and CD27. The different RBD-specific B cell subpopulations were superimposed on total CD19+/CD20+/HLA-DR+ B cells (grey). Legend: IgM+ and naïve IgM+ B cells, blue; IgG+ and memory IgG+ B cells, red; IgA+ and memory IgA+ B cells, orange.

**Supplementary Figure 5.**
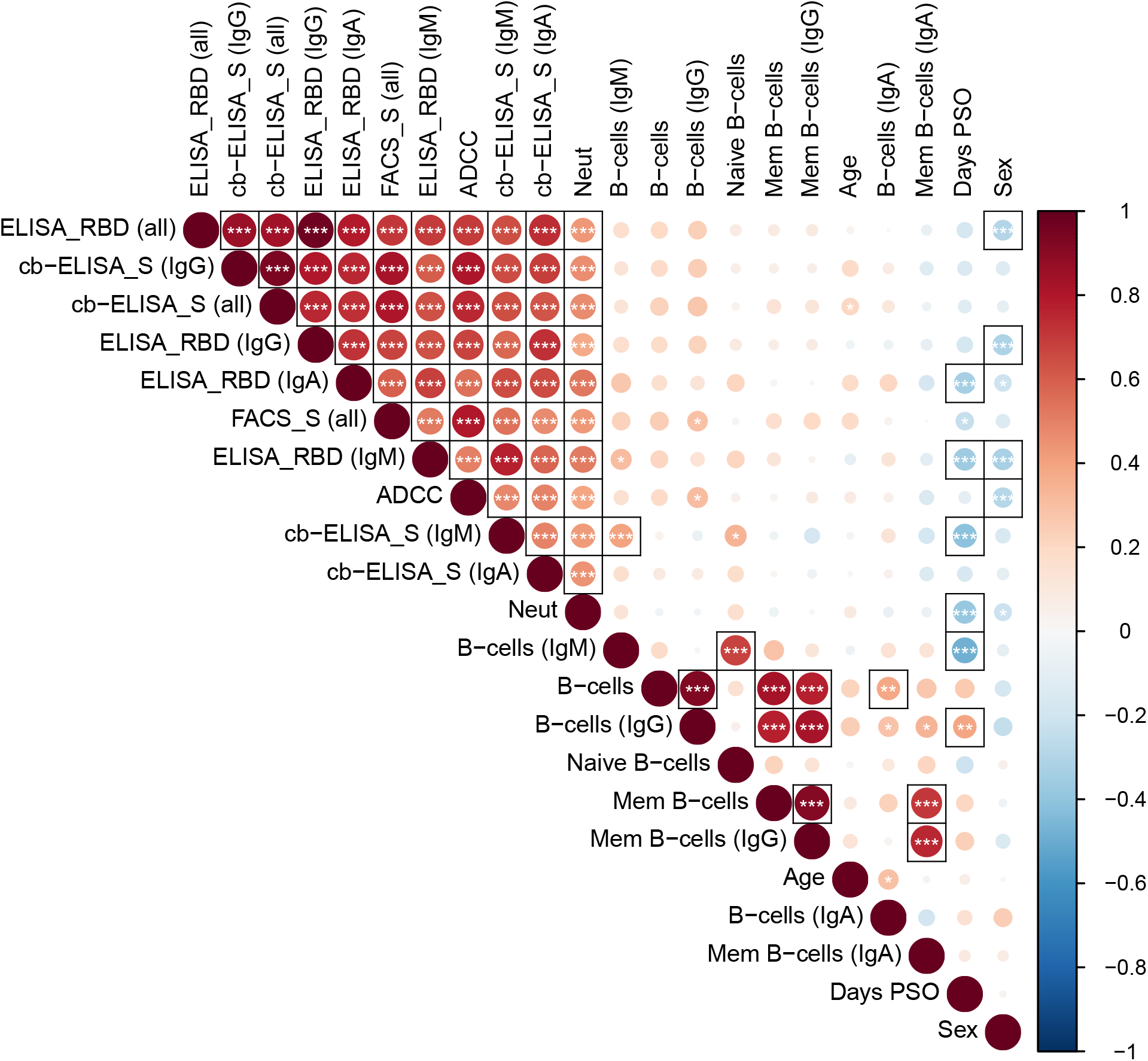
Correlations between serological, immunological and demographic determinants. Correlograms were generated by plotting together all serological, immunological and demographic data obtained from convalescent patients. Circles are color-coded and sized according to the magnitude of the correlation coefficient (r). Red circles represent positive correlations between two variables and blue circles represent negative correlations. Asterisks indicate statistically significant correlations (*P < 0.05, **P < 0.01, ***P < 0.005). Correlation analysis was done using Spearman correlation rank tests. Parameters are clustered hierarchically according to the first principal component (FPC). Black surrounding boxes indicate adjusted p-values < 0.05 using Benjamini-Hochberg multiplicity correction. Legend: Cb-ELISA = cell-based ELISA, Neut = Neutralization, mem = memory, PSO = post-symptom onset, Sex = Female.

